# sAPPα inhibits neurite outgrowth in primary mouse neurons via GABA B Receptor subunit 1a

**DOI:** 10.1101/2025.09.14.675870

**Authors:** Dylan Barber, Casandra Salinas-Salinas, Samah Houmam, Kriti Shukla, Heather C. Rice

## Abstract

Neurite outgrowth is essential for neural circuit formation and is tightly regulated by secreted factors and their receptors. The secreted extracellular domain of the amyloid precursor protein (sAPPα) has been shown to modulate neurite outgrowth. Recently, the gamma amino butyric acid receptor type-B subunit 1a (GABA_B_R1a) was identified as an sAPPα binding partner that mediates its effects on synaptic transmission. Here, we investigated whether this interaction also regulates neurite outgrowth. In primary hippocampal neurons, the GABA_B_R agonist baclofen reduced axon length; whereas, its antagonist CGP54626 increased axon length in primary hippocampal neurons. Moreover, GABA_B_R1a knockout increased axon length and abolished the effect of baclofen. Application of sAPPα reduced axon length, an effect that required the presence of both GABA_B_R1a and the extension domain of sAPPα, which mediates its binding to GABA_B_R1a. Similarly, the APP 17mer peptide, which is sufficient to bind GABA_B_R1a and mimic the effects of sAPP on synaptic transmission, reduced axon outgrowth in wildtype but not in GABA_B_R1a-deficient neurons. Together, these findings indicate that the 1a isoform contributes to GABA_B_R-dependent suppression of neurite outgrowth and mediates the inhibitory effect of sAPPα on neurite outgrowth.

**Statement of Significance:** Amyloid precursor protein (APP) plays a central role in Alzheimer’s disease, yet its normal functions are not fully understood. In this study, we uncover a previously unrecognized role of the GABA B Receptor in mediating the inhibitory effects of sAPPα on neurite outgrowth. These findings provide mechanistic insight into how disruptions in APP signaling could influence both normal brain development and pathological processes in neurodevelopmental disorders and Alzheimer’s disease.

## Introduction

Neurite outgrowth, the process by which neurons extend their axons and dendrites, is essential for establishing neural circuity in the brain. This developmental process is tightly regulated by secreted factors and their receptors. APP is a type I transmembrane protein that undergoes sequential proteolytic processing to generate several fragments, most notably the amyloid-beta (Aβ) peptide which accumulates in Alzheimer’s disease (AD) (Glenner & Wong, 1984). APP processing also leads to the secretion of other physiologically relevant fragments besides Aβ. The initial cleavage of APP by either α-, β-, or η-secretase, releases a large extracellular fragment termed secreted APP (sAPPα, sAPPβ, or sAPPη, respectively (Hefter et al., 2020). During early development, APP expression rises (Hung et al., 1992; Kirazov et al., 2001) and contributes to neurodevelopmental processes such as neurite outgrowth (Chau et al., 2023; Nicolas & Hassan, 2014). Decades of research have shown that primary neurons cultured from APP knock out mice exhibit enhanced neurite outgrowth (Billnitzer et al., 2013; Liu et al., 2021; Perez et al., 1997; Young-Pearse et al., 2008) and also indicates that sAPPα is an important fragment in mediating this function of APP in neurite outgrowth (Billnitzer et al., 2013; Chasseigneaux et al., 2011; Hasebe et al., 2013; Milward et al., 1992, Young-Pearse et al., 2008).

sAPPα interacts with the gamma-aminobutyric acid receptor type B (GABA_B_R) (Dinamarca et al., 2019; Rem et al., 2023; Rice et al., 2019; Schwenk et al., 2016), a metabotropic receptor for the inhibitory neurotransmitter GABA. GABA_B_R is an obligate heterodimer composed of two subunits. Subunit 1 binds GABA; subunit 2 couples intracellularly to Guanine nucleotide-binding (G) proteins (Pin & Bettler, 2016). Subunit 1 exists in two main isoforms, with the 1a isoform containing two additional N-terminal sushi domains that are absent in the 1b isoform (Pin & Bettler, 2016). The extension domain (ExD) within the extracellular region of APP was found to bind specifically to the first sushi domain of the 1a isoform (GABA_B_R1a) (Rice et al., 2019). GABA_B_R1a was shown to mediate effects of sAPPα on synaptic transmission, and a synthetic 17 amino acid peptide within the ExD of sAPPα (APP 17mer) was sufficient to bind GABA_B_R1a and mimic these effects(Rice et al., 2019). However, whether GABA_B_R1a also mediates other functions of sAPPα is not yet known.

The protein expression of both GABA_B_R and APP rise during the first few weeks of rodent development (Fritschy, 1999; Hung et al., 1992; Khoshdel-Sarkarizi et al., 2019; Kirazov et al., 2001; Schwenk et al., 2016). Moreover, both APP and GABA_B_R have been implicated independently in the regulation of similar neurodevelopmental processes, including neurogenesis (Demars et al., 2011; Giachino et al., 2014; Ma et al., 2008), neuronal migration (Bony et al., 2013; Callahan et al., 2017; Rice et al., 2012; Young-Pearse et al., 2007), synaptogenesis (Fiorentino et al., 2009; Meier et al., 2008; Tyan et al., 2012) and neurite outgrowth (Billnitzer et al., 2013; Bony et al., 2013; Chasseigneaux et al., 2011; Favuzzi et al., 2021; Gakhar-Koppole et al., 2008; Hasebe et al., 2013; Hoe et al., 2009; Milward et al., 1992; Osterfield et al., 2008; Perez et al., 1997; Sernagor et al., 2010; Young-Pearse et al., 2008). Here, we sought to determine whether GABA_B_R mediates effects of sAPPα on neurite outgrowth.

Our study demonstrates that knockout of GABA_B_R1a promotes axon outgrowth in primary hippocampal neurons. We also show that both sAPPα and APP 17mer inhibit axon outgrowth and that these effects can be reversed by either removing the ExD of sAPPα or by genetically ablating GABA_B_R1a. Together, these findings indicate that GABA_B_R1a is a key mediator of the inhibitory effect of sAPPα on axon outgrowth.

## Materials and Methods

### Mouse models

All animal procedures were performed in accordance with the [Author University] animal care committee’s regulations. C57BL/6J mice (Jackson Laboratories) were group housed in the AALAS-accredited OMRF vivarium operating on a 14:10 hour light/dark cycle with ad libitum access to food and water. Timed mated females and their embryonic litters used to generate primary neurons for experimentation. GABA_B_R1a knockout (KO) mice generated by the VIB-KU Leuven Center for Brain & Disease Research Mouse Expertise Unit with support from VIB Discovery Sciences. Sperm from GABA_B_R1a KO mice were provided by Joris de Wit, VIB-KU Leuven Center for Brain & Disease Research, Leuven, Belgium and then mice were rederived at the Texas A&M Institute for Genomic Medicine. For the generation of embryonic litters from GABA_B_R1a KO mice, two heterozygous adults were paired for timed mating. The resulting litters contained WT (+/+), Het (+/-), and KO (-/-) embryos, which were used for primary cultures. Genotyping was performed on utilizing the KAPA Hotstart Mouse Genotyping Kit (Kapa Biosystems KK7352) with the following primers: mGABBR1Fwd1 [5’-GGAAGAAGAACAGGGGGA-3’], mGABBR1Rev1 [5’-AGGAGGTCAGGAGTTGTG-3’].

### Primary mouse hippocampal neuron culture

Hippocampal neurons were isolated from embryonic day 18 (E18) mouse brains. Hippocampi were dissected by decapitating the embryos and placing the heads into ice-cold Hank’s Balanced Salt Solution without magnesium and calcium (Gibco 14-175-095), supplemented with 2.5mM HEPES, 30mM D-glucose, 1mM CaCl_2_, 1mM MgSO_4_, and 4mM NaHCO3 (cHBSS). The brain was extracted, and the hippocampus was dissected into ice-cold cHBSS. Tissues were dissociated by incubation in cHBSS with 0.25% trypsin (Gibco 15090046) and DNAse [50µg/mL] (Sigma-Aldrich 11284932001) for 15 minutes at 37 °C, followed by gentle trituration using a flame-polished Pasteur pipette. Cells were washed with cHBSS 3 times and plated at 75,000 cells/mL on 12mm coverslips precoated with poly-D-lysine (Neuvitro GG-12-PDL), coated with laminin [0.001mg/mL] (Invitrogen 114956-81-9 L2020). Coverslips were placed into a 12-well cell culture dish (Corning 353043) filled with cHBSS. Neurons were fed by half media change after 3 hours with Hippocampal Feeding Media (HFM): Neurobasal medium (Invitrogen 21103049) supplemented with 1X B27 (Gibco 17504044), 0.25X GlutaMAX (Gibco 35050061), 0.24% D-glucose (Thermo Fisher Scientific A16828.0C), penicillin/streptomycin [20U/mL] (Invitrogen 15140122) and 24.2µM β-mercaptoethanol (Thermo Fisher Scientific 2198502). For the litters from the GABA_B_R1a^+/-^ matings, embryonic tissue samples were used for genotyping as described above. The dissected hippocampi were dissociated separately and plated onto batches of coverslips independently. After same-day genotyping, coverslips of the three possible genotypes GABA_B_R1a^+/+^, GABA_B_R1a^+/-^, and GABA_B_R1a^-/-^ were selected for treatment. Cells were treated with purified proteins, synthetic peptides, or pharmacological agents by bath application 3 hours after plating. Purified sAPPα and sAPPαΔExD proteins (see plasmids and purification methods below) or synthetic APP 17mer and scrambled 17mer peprides (described below) were applied at a final concetration of 500nM. The GABA_B_R agonist baclofen (Sigma Aldrich 63701-55-3) and antagonist (Sigma Aldrich SML3136) were both bath applied at 10µM. Neurons were maintained in HFM and fixed 72 hours after plating.

### Plasmids

sAPP-Fc constructs were provided by Dr. Joris de Wit, VIB-KU Leuven Center for Brain & Disease Research, Leuven, Belgium. sAPP-Fc constructs were originally generated by PCR-amplifying the following regions of mouse APP695: sAPPα= 18-612aa; sAPPαΔExD= 19-194aa & 228-596aa. Each of the PCR fragments were subcloned between and in frame with the prolactin signal peptide and human Fc in the pLP-FLAG-IgG vector using Gibson Assembly (NEB).

### Protein Purification

Secreted C-terminally Fc-tagged proteins were expressed by transient transfection using polyethylenimine (PEI) (Kyfora Bio 23966) in HEK293T cells and collected in serum-free Opti-MEM (Gibco 31985088). Conditioned medium was passed through a Protein-G Sepharose packed column (Cytiva 17061802) at 4 °C, washed with 250 mL wash buffer (50mM Tris pH 8.0, 450 mM NaCl, 1 mM EDTA), and Fc tag cleaved O/N with GST-tagged 3C PreScission Protease (Cytiva 27084301) in cleavage buffer (50mM Tris pH 8.0, 150 mM NaCl, 1 mM EDTA, 1 mM DTT). Cleaved protein was collected in the eluate and the protease separated from the eluted proteins using a Glutathione Sepharose (Cytiva 17513201) column collecting in TNE buffer (50mM Tris pH 8.0, 150 mM NaCl, 1 mM EDTA). Proteins were dialyzed against Phosphate-Buffered Saline (PBS) O/N, concentrated using centrifugal filter units (Millipore Sigma UFC9010), and depleted of endotoxin with Pierce™ High Capacity Endotoxin Removal Spin Columns (Thermo Scientific 88274). Protein concentration was determined by BCA Protein Assay (Thermo Scientific 23227) and verified by Coomassie SDS-PAGE.

### Synthetic peptides

The following peptides were synthesized by Insight Biotechnology at >98% purity: APP 17mer (204-220AA of APP695): acetyl-DDSDVWWGGADTDYADG-amide Scrambled 17mer: acetyl-DWGADTVSGDGYDAWDD-amide

### Immunocytochemistry and light microscopy

For fluorescence staining, cells were fixed with 4% PFA in phosphate buffed saline (PBS) for 15 minutes. Fixed samples were washed three times in PBS, five minutes per wash. The fixed neurons were then blocked with 3% bovine serum albumin in PBS supplemented with TritonX100 (0.2%) O/N at 4C and stained with primary mouse monoclonal Tau-1 antibody to an axon specific microtubule associated protein (1:1000 dilution; Millipore Sigma MAB3420), primary chicken MAP2 antibody to stain dendrites (1:1000; Abcam Ab5392), and a primary rabbit Tuj1 antibody against pan-neuronal β3 tubulin (1:1000 dilution; Abcam Ab18207). Samples were then secondary antibody conjugated to Alexa Fluor 488 (Southern Bio-tech 6410-30), Alexa Fluor 568 (ThermoFisher A78950), or Alexa Fluor 647 (Biolegend 406414) reconstituted at 1,2, 0.5 mg/mL respectively all used at 1:200 dilution. Light microscopy imaging was performed using an Axioscan7 Whole Slide Scanner at 40X air magnification with the OMRF Imaging Core Facility.

### Image analysis

All analysis was performed blinded to treatment conditions. Blinding was achieved through basic Caesar cipher of files names and pseudo labeling of slides prior to imaging on the Axioscan 7. The region of interest for analysis was 21mm^2^ and neurons were measured within the ROI in clockwise manner starting from a random corner of the image until the target number of neurons were measured (60-90/coverslip). Measurements were taken using ImageJ Simple Neurite Tracer (SNT) software. Neurite lengths were measured from the axon hillock to the tip of the longest primary neurite stained by Tau-1 (Axon Marker). Neurite lengths were determined in three independent experiments using images acquired at 25-40× magnification stained by Tau-1 (Axon Marker) and MAP2 (Dendrite Marker) for the purpose of differentiating the neuronal compartments. Images were imported into ImageJ using the bio-formats importer plug-in. The images were then imported into the SNT plug-in and axon filaments were traced using the filament tracer algorithm. Blinding was removed after datasets were collected.

### Statistical analysis

Statistical analysis (analysis of variance) was performed using Graphpad Prism (Boston, MA) software. Normality was determined by Anderson-Darling and Shapiro-Wilk test. Significance was determined using Kruskal-Wallis with Dunn’s multiple comparison post hoc test. Graphs were produced using Graphpad Prism.

## Results

GABA_B_R has been implicated in the regulation of neurite outgrowth of mouse primary neurons (Bony et al., 2013). To confirm these findings, we treated hippocampal neurons isolated from wildtype C57BL/6J E18 mouse embryos with baclofen, an agonist of GABA_B_R, and CGP54626, an antagonist of GABA_B_R. Neurons were treated 3 hours after plating and immunostained after 3 days in vitro (3 DIV) for MAP2 to label dendrites and Tau to label axons (Figure 1A). Consistent with previous findings (Bony et al., 2013), we found that bath application of 10µM baclofen significantly reduced axon length by 9% compared to untreated controls and 10µM CGP 54626 increased axon length by 14% (Figure 1B). Baclofen and CGP54626 are not isoform-specific in their modulation of the GABA_B_R; therefore, to determine whether GABA_B_R1a, which specifically binds sAPPα, regulates neurite outgrowth, we performed neurite outgrowth assays on primary neurons isolated from littermates that were full KO (GABA_B_R1a^-/-^), heterozygous KO (GABA_B_R1a^+/-^), or wildtype (GABA_B_R1a^+/+^) (Figure 1C). We found that axon length significantly increased by 27% in GABA_B_R1a^+/-^ neurons and 9% in GABA_B_R1a^-/-^ as compared to GABA_B_R1a^+/+^. (Figure 1D). To determine the contribution of GABA_B_R1a to the reduction in axonal length by baclofen, we treated GABA_B_R1a^-/-^, GABA_B_R1a^-/+^ and GABA_B_R1a^+/+^neurons with 10µM baclofen. Baclofen significantly reduced axon length by 40% in both wildtype and heterozygous GABA_B_R1a KO neurons (Figure 1E). However, baclofen had no significant effect on axon length in homozygous GABA_B_R1a KO neurons (Figure 1E). These findings demonstrate that GABA_B_R1a reduces axon outgrowth in primary neuron cultures.

**Figure 1:**
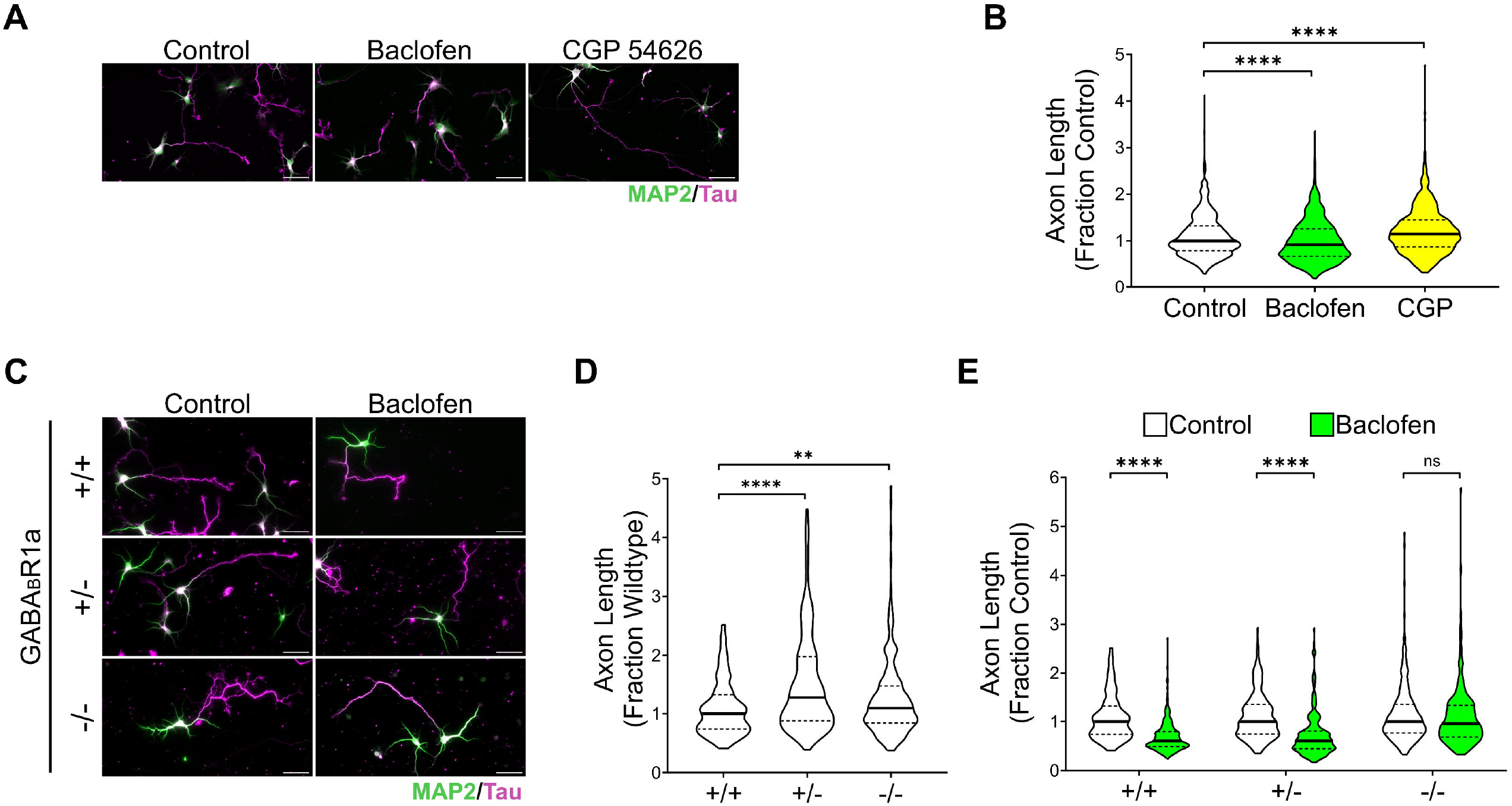
Modulation of GABA_B_R influences axon outgrowth in primary neurons. **A)** Representative images of hippocampal primary neurons derived from C57BL/6J E18 mouse pups treated at DIV 0 with baclofen (GABA_B_R agonist) or CGP54626 (GABA_B_R antagonist) and immunostained at DIV3 with MAP2 (green, dendritic marker) and Tau (magenta, axonal marker). **B)** Baclofen treatment significantly decreased axon length (N = 180-220 neurons/trial across 5 trials; Median = 0.9187, IQR = 0.6672-1.259). CGP54626 treatment significantly increased axon outgrowth (N = 180-220 neurons/trial across 5 trials; Median 1.145 = IQR 0.8696-1.451) compared to untreated controls. **C)** Representative images of hippocampal primary neurons derived from wildtype (+/+), heterozygous GABA_B_R1a KO (-/+), and homozygous GABA_B_R1a KO (-/-) E18 littermates treated at DIV 0 with baclofen and immunostained at DIV3 with MAP2 (green, dendritic marker) and Tau (magenta, axonal marker). **D)** Axon length was significantly increased in both GABA_B_R1a (-/-) and (-/+) neurons compared to controls (N = 60-90 neurons/trial across 3 trials; Medians = 1.096 and 1.274; IQR = 0.8413-1.469 and 0.8797-1.977, respectively). **E)** Baclofen treatment significantly decreased axon length in primary neurons derived from GABA_B_R1a (+/+) and (-/+) mice, (N = 60-90 neurons/trial across 3 trials; Medians = 0.6067 and 0.6067, IQR = 0.4911-0.7975 and 0.4461-0.8122, respectively). Controls from panel D and E are the same, normalized differently for comparison. Graphs show medians and interquartile ranges. Scale bars, 50µm. Kruskal-Wallis with Dunn’s multiple comparison post hoc test were used. **P <0.01, ****P<0.0001; ns, not significant (P >0.05)

sAPPα binds to the sushi-1 domain present in the 1a isoform of GABA_B_R, (Rice et al., 2019); therefore, we sought to determine whether GABA_B_R1a mediates the effects of sAPPα on neurite outgrowth. Since sAPPα binds the sushi-1 domain at a dissociation constant (KD) of 431 nM (Rice et al., 2019), we treated wildtype mouse hippocampal neurons with 500 nM sAPPα and measured axon length after 3 DIV. sAPPα (Figure 1A) was affinity purified from transfected–human embryonic kidney (HEK) 293T cell supernatants (Figure 2B). Treatment with 500 nM sAPPα reduced axon length by 36% compared to untreated controls (Figure 2C,D). To determine if the GABA_B_R1a binding region is required for this effect, we also treated wildtype neurons with an sAPPα lacking the ExD (sAPPα-ΔExD) (Figure 2B). Treatment with 500 nM sAPPα-ΔExD did not significantly affect axon length compared to untreated controls (Figure 2C,D), indicating that inhibition of axon outgrowth by sAPPα requires the ExD. Previously, binding between sAPPα and GABA_B_R1a was mapped to a 17aa region within the ExD of APP (Rice et al., 2019). This APP 17mer was sufficient to mimic the GABA_B_R1a-dependent effects on synaptic transmission (Rice et al., 2019). To determine if APP 17mer could also modulate axon outgrowth, we applied 500nM APP 17mer or its scrambled control. APP 17mer reduced axon length by by 11% compared to untreated controls (Figure 2C,D); whereas, a scrambled 17mer peptide did not significantly affect axon length. Together, these findings are consistent with a role for GABA_B_R1a in mediating the inhibitory effect of sAPPα on neurite outgrowth in primary mouse neurons.

**Figure 2:**
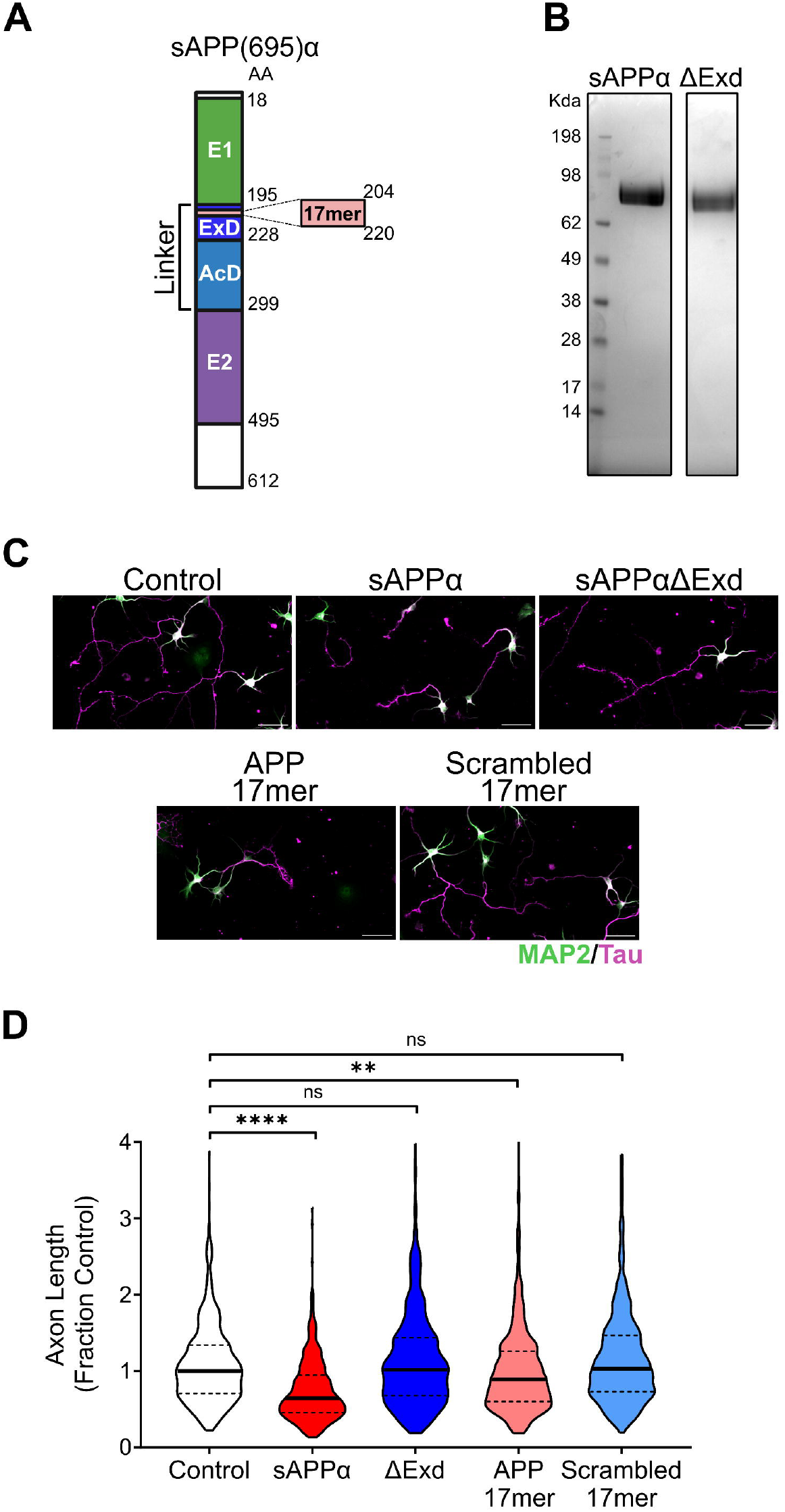
sAPPα reduces axon length in primary neurons. **A)** Cartoon of the domains of sAPP(695)α. **B)** Coomassie stain of sAPPα and sAPPα-ΔExD purified proteins. **C)** Representative images of hippocampal primary neurons derived from C57BL/6J E18 mouse pups treated at DIV 0 with sAPPα, sAPPα-ΔExD, APP 17mer, or scrambled 17mer and immunostained at DIV3 with MAP2 (green, dendritic marker) and Tau (magenta, axonal marker). **D)** Treatment with sAPPα and 17mer decreased axon length in primary neurons compared to untreated controls (N = 180-200 neurons/trial across 3 trials; Medians = 0.6461 and 0.8894, IQR = 0.4562-0.9462 and 0.6007-1.258, respectively). Treatment with sAPPα-ΔExD and scrambled 17mer had no significant effect on axon length. Graphs show medians and interquartile ranges. Scale bars, 50µm. Kruskal-Wallis with Dunn’s multiple comparison post hoc test were used. **P <0.01, ****P<0.0001; ns, not significant

To test whether GABA_B_R1a is required for the inhibition of axon outgrowth by sAPPα, we cultured primary neurons from GABA_B_R1a^-/-^, GABA_B_R1a^+/-^, or GABA_B_R1a^+/+^ littermates and treated each genotype with 500nM sAPPα, sAPPα-ΔExD, APP 17mer, and scrambled 17mer. sAPPα reduced axon length by 23% in GABA_B_R1a^+/+^ neurons and 42% in GABA_B_R1a^+/-^ neurons; whereas, sAPPα-ΔExD had no significant effect on axon length (Figure 3A,B). Similarly, 500 nM APP 17mer resulted in a 34% reduction in axon length of GABA_B_R1a^+/+^ and 36% reduction in GABA_B_R1a^+/-^ neurons compared to untreated controls; whereas, scrambled 17mer had no significant effect on axon length (Figure 3A,B). Strikingly, the effects of sAPPα and APP 17mer were abolished in GABA_B_R1a^-/-^ neurons (Figure 3A,B). Together, these findings demonstrate that sAPPα requires the presence of GABA_B_R1a to inhibit neurite outgrowth.

**Figure 3:**
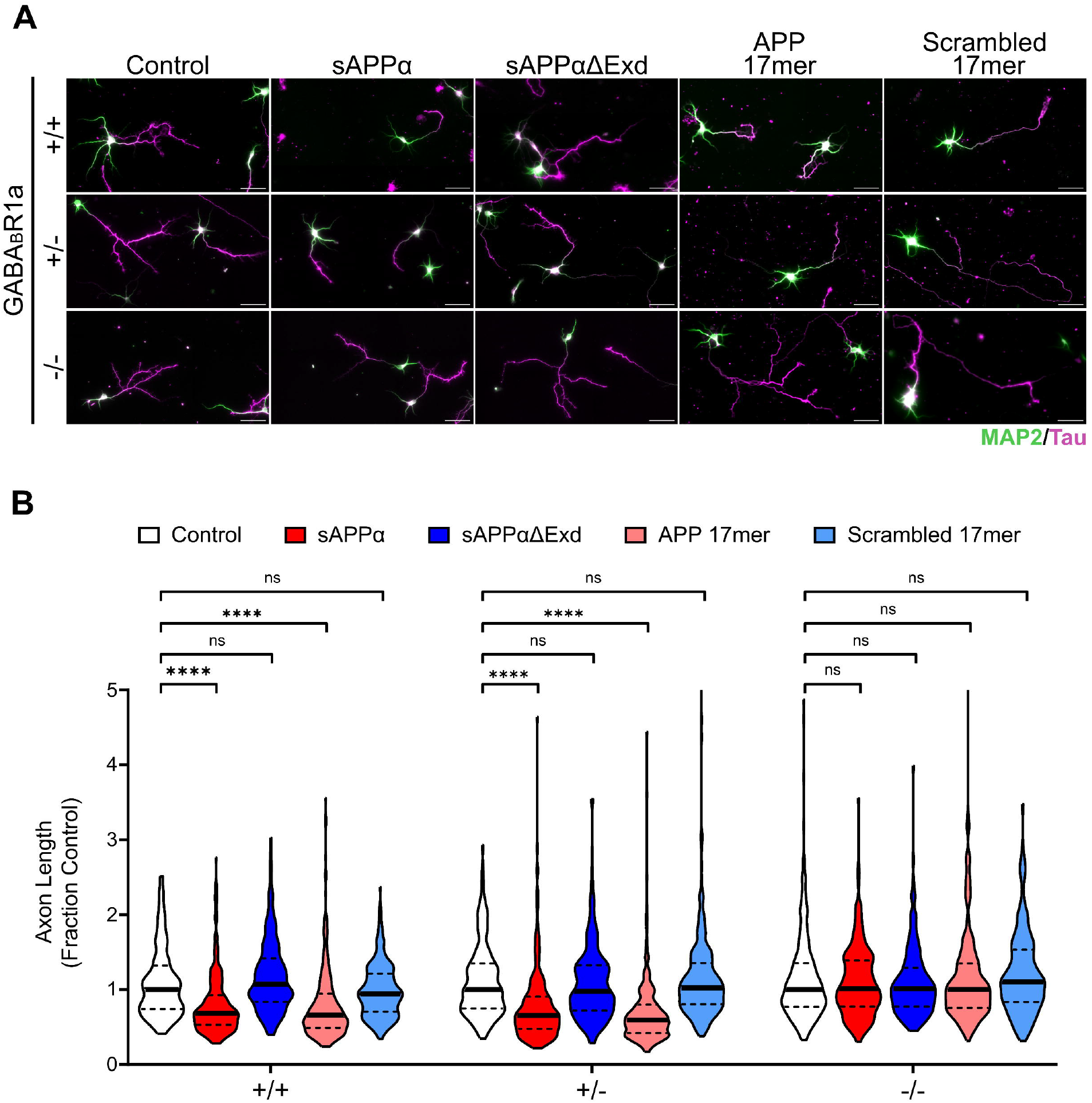
The effects of sAPPα and APP 17mer are abolished in GABA_B_R1a^-/-^ neurons **A)** Representative images of hippocampal primary neurons derived from wildtype (+/+), heterozygous GABA_B_R1a KO (-/+), and homozygous GABA_B_R1a KO (-/-) E18 littermates treated at DIV 0 with sAPPα, sAPPα-ΔExD, APP 17mer, or scrambled 17mer and immunostained at DIV3 with MAP2 (green, dendritic marker) and Tau (magenta, axonal marker). **B)** Treatment with sAPPα significantly reduced axon length in both GABA_B_R1a +/+ and -/+ neurons (N = 60-90 neurons/trial across 3 trials; Medians = 0.6613 and 0.5940, IQR = 0.5305-0.9274 and 0.4785-0.9049, respectively). Treatment with APP 17mer significantly reduced axon length in both GABA_B_R1a +/+ and -/+ neurons (N = 60-90 neurons/trial across 3 trials; Medians = 0.6832 and 0.6568, IQR = 0.4906-0.9480 and 0.4213-0.7989, respectively). Treatment with sAPPα and APP 17mer had no significant effect on axon length in GABA_B_R1a -/- neurons. Untreated controls in 3B are the same neurons as in Figures 1 D,E. Graphs show medians and interquartile ranges. Scale bars, 50µm. Kruskal-Wallis with Dunn’s multiple comparison post hoc test were used. ****P<0.0001; ns, not significant

## Discussion

Here, we found that 500nM sAPPα reduces axon outgrowth in mouse primary hippocampal neurons (Figure 2D). This effect required the presence of both GABA_B_R1a and the ExD of sAPPα, which mediates its interaction with GABA_B_R1a (Figure 3A-B). The APP 17mer, which was previously shown to bind GABA_B_R1a and mimic the effects of sAPP on synaptic transmission(Rice et al., 2019), also reduced axon outgrowth in WT but not in GABA_B_R1a KO neurons (Figure 3A-B). Together, these findings demonstrate that sAPPα inhibits neurite outgrowth in primary mouse neurons via GABA_B_R1a.

Previously, GABA_B_R has been implicated in regulating neurite outgrowth, with activation by the agonist baclofen reported to reduce axon length and inhibition by CGP reported to enhance axon length in primary neurons (Bony et al., 2013). Our present study not only confirms these findings (Figure 1B) but also establishes a role for the GABA_B_R1a isoform in this effect. We show that both partial and full ablation of GABA_B_R1a promotes neurite outgrowth (Figure 1D) and the inhibition of neurite outgrowth by the GABA_B_R agonist baclofen requires the presence of GABA_B_R1a (Figure 1E). Thus, the 1a isoform contributes to GABA_B_R-dependent suppression of neurite outgrowth.

Decades of research across multiple groups has demonstrated that both full length APP and sAPPα play a role in neurite outgrowth (Chau et al., 2023). The prevailing consensus is that ablation of full-length APP promotes neurite outgrowth in primary neuron cultures (Billnitzer et al., 2013; Liu et al., 2021; Perez et al., 1997; Young-Pearse et al., 2008). Paradoxically, numerous studies report that application of sAPPα also promotes neurite outgrowth (Billnitzer et al., 2013; Chasseigneaux et al., 2011; Hasebe et al., 2013; Milward et al., 1992; Young-Pearse et al., 2008). As an exception, one study found that neurons co-cultured with APP KO astrocytes had elongated neurites, suggesting that sAPPα can, under certain conditions, inhibit neurite outgrowth (Perez et al., 1997). Consistent with this and aligning with the observation that APP ablation enhances neurite outgrowth, our current findings indicate that sAPPα can reduce neurite outgrowth. We speculate the directionality of the effect of sAPPα may depend on its context-dependent interactions with different binding partners. Since sAPPα binds the sushi-1 domain at a KD of 431 nM (Rice et al., 2019), we treated neurons with 500 nM sAPPα. Previous studies applied sAPPα at 1-150nM (Billnitzer et al., 2013; Hasebe et al., 2013; Young-Pearse et al., 2008) which may be insufficient to bind GABA_B_R1a. Instead, at lower concentrations, binding to other interactors, such as Integrin β1 (Young-Pearse et al., 2008) may promote neurite outgrowth. Interestingly, synaptic activity is known to regulate APP processing (Cirrito et al., 2005; Tampellini et al., 2009) and could thereby tightly regulate neurite outgrowth.

APP and GABA_B_R have each been independently implicated in neurodevelopmental disorders, including fragile X syndrome (Henderson et al., 2012; Kang et al., 2017; Ray et al., 2016; Westmark, Chuang, et al., 2016; Westmark et al., 2011; Westmark, Sokol, et al., 2016), autism spectrum disorders (Fatemi et al., 2009; Frackowiak et al., 2014, 2020; Li et al., 2022; Wegiel et al., 2012), and Down syndrome (Best et al., 2007; Cheon et al., 2008; Kleschevnikov, Belichenko, Gall, et al., 2012). In Down syndrome, APP expression is increased due to triplication (Cheon et al., 2008), and pharmacological inhibition of GABA_B_R has been shown to rescue deficits in a Down syndrome mouse model (Kleschevnikov, Belichenko, Faizi, et al., 2012). Our findings support a functional link between dysregulation of APP and GABA_B_R signaling pathways in neurodevelopment. The ability of a short APP-derived 17mer peptide to activate this pathway highlights the therapeutic potential of targeting APP-GABA_B_R1a signaling for the treatment of neurodevelopmental disorders.

## Acknowledgements

We thank Yvonne Thomason, Suyesha Bhandari, and Charles Lacy for technical support and Dominika Siodlak for scientific discussions. Image acquisition was supported by the OMRF Imaging Core Facility and the OUHSC Molecular Analysis and Cellular Imaging Core of the “Cellular and Molecular GeroScience CoBRE”. Data processing and analysis were supported by the OMRF Center for Biomedical Data Sciences.

## Notes

**Conflict of Interest:** No

**Funding Sources:** This work was supported by the National Institutes of Health (R35GM142726, T32AG052363, P20GM1125528), the Presbyterian Health Foundation (Pilot Research Funding) and the OUHSC College of Medicine Alumni Association (Pilot Research Funding).

### Competing Interest Statement

The authors have declared no competing interest.

